# Topology Determines DNA Origami Diffusion in Intestinal Mucus

**DOI:** 10.1101/2025.04.25.650571

**Authors:** Matteo Tollemeto, Lars J.M.M. Paffen, Lasse H. E. Thamdrup, Anja Boisen, Jan van Hest, Tania Patiño Padial

**Author notes:** **Corresponding authors:** Tania Patiño Padial.

## Abstract

Efficient nanomedicine delivery across mucosal barriers remains a challenge, due to the complex and poorly understood relationship between nanoparticle design and mucus transport. Here, we present DNA origami as a platform to investigate how nanoparticle shape and ligand patterning influence diffusivity in mucus. By decoupling these parameters while maintaining identical material composition, we systematically evaluated the diffusion of rod, icosahedral, and rectangular nanostructures using high-resolution single-particle tracking. Our results reveal that diffusivity in mucus is not solely determined by shape or functionalization alone, but by their interplay: while unmodified rods diffused poorly, their mobility increased significantly upon antibody functionalization, reaching a maximum at an intermediate ligand density. In contrast, rods and icosahedra exhibited less pronounced and non-optimal responses to surface modification. These findings highlight the importance of topology-specific optimization in nanoparticle design and demonstrate the utility of DNA nanotechnology to uncover design rules for next generation mucus-penetrating drug delivery systems.

## Introduction

Nanomedicines have emerged as a promising strategy for enhancing the bioavailability of drugs by improving their stability, solubility, and accumulation at specific target locations. However, biological barriers continue to present major challenges for the efficient and targeted delivery of nanoparticles.^1–3^ One important barrier is the mucus layer that lines various epithelial surfaces, including those in the respiratory and gastrointestinal tracts.^4^ Mucus is a complex biological medium composed of water, mucins (glycoproteins), DNA, proteins, lipids and cell debris, and its primary function is to lubricate and serve as a protective shield to exposed epithelia. The natural protective role of mucus presents a significant obstacle to nanomedicine transport due to its complex structure and high viscosity. In this regard, not only does the mucus barrier act as a physical impediment, but it also influences the diffusion and absorption of nanoparticles, ultimately affecting therapeutic outcomes.^5,6^While there are numerous examples of mucus-penetrating nanoparticles for drug delivery,^7,8^ the mechanisms governing nanoparticle behaviour within mucus under varying physiological and pathophysiological conditions remains insufficiently understood. Several studies have shown that factors such as size^9^, shape^10^, surface properties^11,12^, and rigidity^13^ significantly affect nanoparticle diffusion through intestinal mucus. Despite the rapid and exciting advancements in the field, a main challenge towards robust mechanistic understanding of nanoparticle behaviour in mucus relies on the difficulty to control multiple nanoparticle topological features, including shape, size and ligand functionalization while keeping the same material properties.

In this context, DNA origami allows for the design and assembly of 1D, 2D^14^ and 3D^15–17^ nanostructures with full control over size and shape. This can be achieved by the folding of a ssDNA scaffold using shorter compatible DNA strands (staples). Due to its unique programmability and design tunability while keeping the same material properties, DNA origami is emerging as a valuable platform for studying and optimizing nanoparticle interactions with biological systems. ^18–27^

Here, we aimed at exploring DNA origami as a model to investigate how topological features determine nanoparticle diffusion in intestinal mucus. For this, we employed three distinct DNA-origami shapes, i.e. icosahedron, rectangle and rod, and we functionalized them with the therapeutic anti-EGFR (Epidermal Growth Factor Receptor) antibody, given its potential for colorectal cancer applications.^28–30^ To investigate the diffusion dynamics of DNA origami within mucus in space and time, we employed single particle tracking (SPT), a high-resolution optical microscopy technique that allows for the tracking of individual nanostructures, including DNA origami^31,32^ with unique spatiotemporal resolution.

By harnessing the precise and programmable design features of DNA origami, we provide a systematic approach to understanding how nanostructure design impacts mucus penetration. This study offers insights that could aid in the rational design of more effective drug delivery systems, contributing to improved bioavailability and therapeutic efficacy.

## Results

### DNA origami design, assembly and characterization

One commonly recognized criterion for achieving optimal diffusion across mucus layers is the size of the nanoparticles, with those below 200 nm generally considered most effective.^33,34^ This size range allows nanoparticles to navigate through the dense meshwork of mucin present in mucus, enhancing their ability to penetrate mucosal barriers. Therefore, when designing the shapes and particles for our study, we specifically selected DNA origami structures that align with this criterion, where all DNA origami structures (i.e. rods, rectangles and icosahedrons) were below the 200 nm threshold. These nanostructures were constructed through bottom-up self-assembly, involving the folding of a long single-stranded DNA (ssDNA) scaffold. For the rods, the scaffold utilized was p7560, while for rectangles and icosahedra, the M13mp18 scaffold was employed. The assembly process was achieved by employing multiple ssDNA strands complementary to specific regions of the scaffold (i.e. ssDNA staple strands). Both ssDNA scaffold and strands were mixed and subjected to thermal annealing, allowing the nanostructures to self-assemble through complementary base pairing interactions (**Figure 1A**). Gel electrophoresis was utilized to confirm the self-assembly of the scaffold into the three different shapes, where it was observed how the folded structures migrate differently compared to the scaffold. The folded DNA origami nanostructures were purified to remove unbound staples **(Figure 1B)** using a PEG-precipitation approach and characterized by dynamic light scattering (DLS). The rod, rectangle and icosahedron designs displayed hydrodynamic diameters of 161.8 ± 6.1 nm, 92.7 ± 5.6 nm, 95.1 ± 4.8 nm respectively, with a PDI of 0.3 for the rods, 0.19 for the rectangles and 0.18 for the icosahedra (**Figure SI 1**). To further confirm the size and shape of the folded DNA origami nanostructures, we performed AFM imaging. The average size was calculated by measuring 50 DNA origami for each particle shape, where a size of 142.6 ± 2.7 and 23.5 ± 2.4 (height x diameter) for the rods, 94.2 ± 11.7 and 66.8 ± 4.0 (height x length) for the rectangles and 72.7 ± 7.2 nm in diameter for the icosahedra was observed (**Figure 1C**). These measurements are consistent with literature values, indicating dimensions of approximately 140-150 × 20-30 nm for the rods, 90 nm x 70 nm x 2 nm for the rectangles, and 50 nm diameter for the icosahedrons.^35–38^

**Figure 1.**
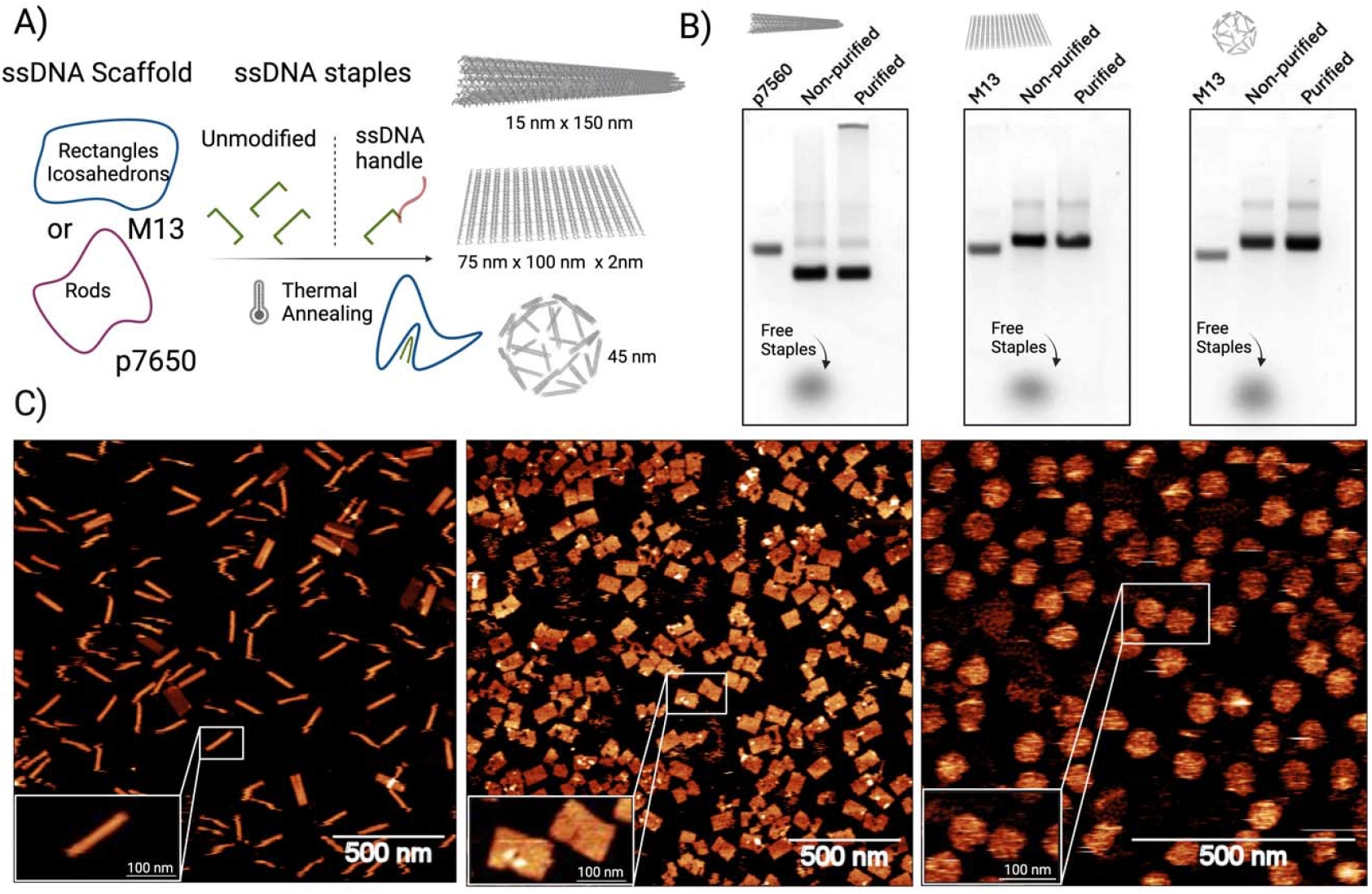
DNA origami assembly. A) Illustrative diagrams depicting the assembly of DNA origami into various shapes, with functionalization at different protein densities. Gel electrophoresis of the three DNA origami shapes displays the scaffold, the non-purified assembled structures, and the purified structures. B) AFM images highlighting the structural details of the various shapes.

### DNA origami diffusion across mucus: the effect of shape

To investigate the impact of DNA origami shape on its diffusion across mucus, the three designs were evaluated in *ex vivo* porcine intestinal mucus using single particle tracking (SPT) under a dSTORM microscope. SPT is a microscopy-based technique that enables a real-time observation of individual particle movements at a high spatial and temporal resolution, making it well-suited for studying diffusion behaviour in complex biological environments. ^27^ For visualization under dSTORM, DNA origami were labeled with the fluorophore ATTO 647. To follow their trajectories, we recorded videos during 10 s at 37°C at 100 FPS. (**Figure 2A**). Representative trajectories are depicted in **Figure 2B**, where we employed a colour code to visually indicate the diffusion coefficient of each trajectory overtime. From these images, clear differences can already be observed among DNA origami shapes, where origami rods displayed trajectories with lower diffusion coefficient values, compared to rectangles and icosahedrons. This effect is more visible in **Figure 2C**, where we extracted and graphically represented different examples of trajectories from each shape, suggesting that the DNA origami nanorods remain largely confined within a specific region of the mucus, whereas both rectangles and icosahedron display higher area exploration. From the SPT of randomly selected particles (N=50), we calculated their mean squared displacement (MSD) (**Figure 2D**) as previously reported.^39^ The MSD of particles serves as an indicator of their diffusive behavior. In this regard, the MSD exponent (α), obtained by fitting MSD as a function of time (⟨*r*2(*t*)⟩∼*t*^*α*^) determines whether the particles exhibit normal diffusion (*α*≈1), subdiffusion (*α*<1), or superdiffusion/ballistic motion (*α*>1).^40^ Interestingly, while both rectangles and icosahedrons displayed linear MSDs with an value close to 1 (0.98 ± 0.02 and 1.037 ± 0.01, respectively), indicating Brownian diffusion, rods exhibited a slight subdiffusive behavior with an α value less than 1 (0.92 ± 0.02). This subdiffusion in rods suggests hindered translational motion, likely due to rotational constraints or interactions with the surrounding environment, which could be explained by their higher anisotropy compared to rectangles and icosahedrons. From the MSDs, we calculated the diffusion coefficients, where significant differences were observed (p < 0.05) among the three shapes, with values of 0.23 ± 0.01 μm^2^/s for rods, 0.44 ± 0.02 μm^2^/s for rectangles, and 0.75 ± 0.02 μm^2^/s for icosahedra (mean ± SEM) (**Figure 2E**). It is noteworthy that despite DNA nanorods and icosahedrons having a comparable theoretical volume (See Supporting Table 1), their diffusion coefficient in mucus is significantly different. This can be explained by the observed differences in the hydrodynamic radius and diffusion behaviour. This can be also observed for rectangles, whereas despite having a significant lower volume compared to icosahedrons, their diffusivity is not higher. We attribute this effect to the fact that both rectangles and rods have a higher anisotropy, resulting in higher hydrodynamic radius and constrained translational diffusivity. Overall, these results highlight the need to consider not only the theoretical particle sizes and volumes but also shape and anisotropy effects on the hydrodynamic size when studying nanoparticle diffusivity within mucus.

**Figure 2.**
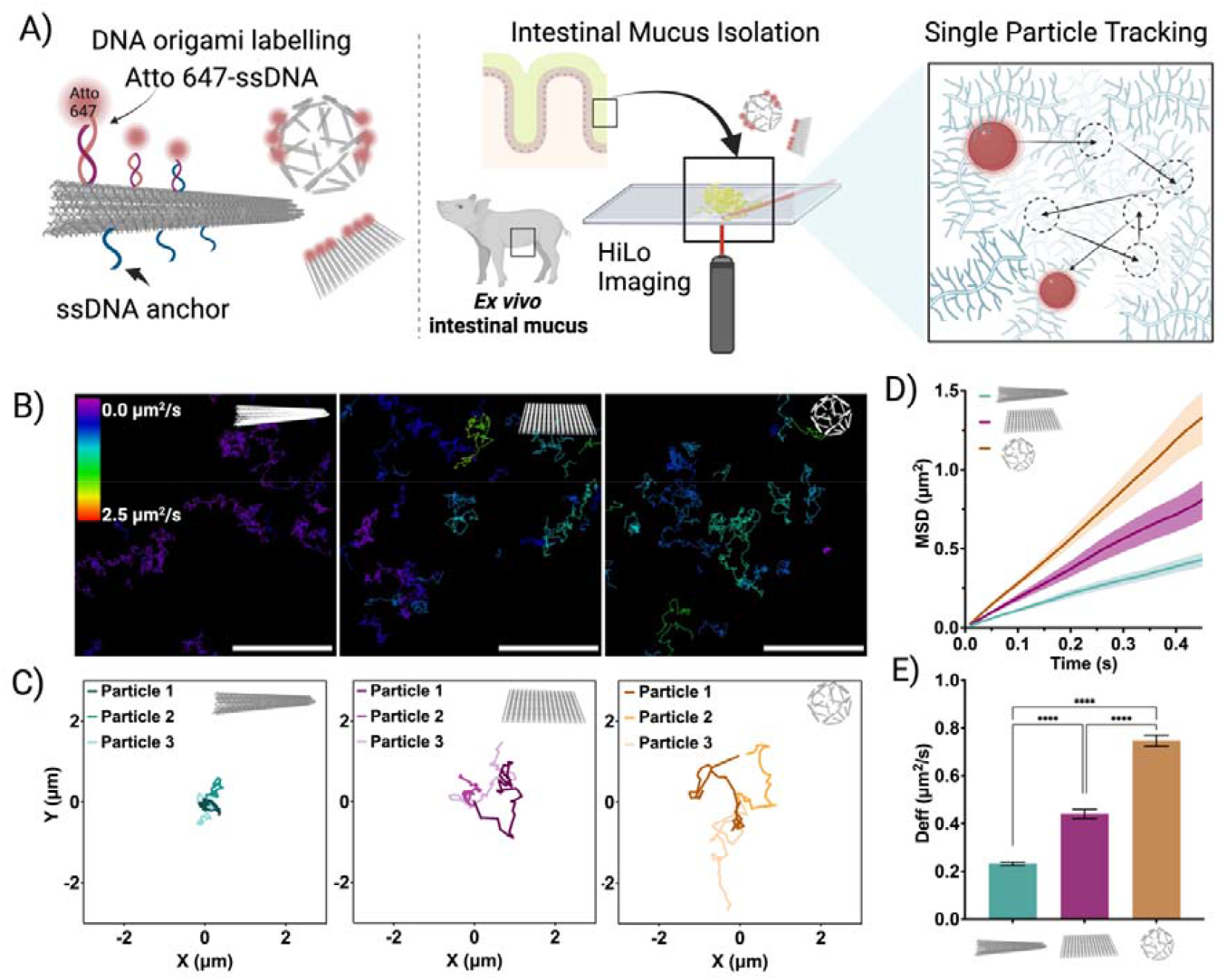
DNA origami tracking. A) Illustrative diagrams showing the tracking process of Atto 647-labeled DNA origami. Initially, *ex vivo* intestinal mucus is extracted from healthy piglets and incubated with DNA origami. The particles are then tracked using HiLo microscopy. B) Top: Color-coded diffusion tracks for different shapes (scale bar 5 μm) C) Three representative tracks for each shape. D) Mean square displacement (MSD) analysis for different shapes in mucus. E) Diffusion coefficients extracted from the MSD analysis for the various shapes. Results are shown as the mean ± standard error of the mean (N=50). Asterisks ‘****’ denote statistically significant differences between groups, with p ≤ 0.0001.

Interestingly, our findings partially contrast with previous studies by Bao et al. and Yu et al., which demonstrated that rod-shaped particles generally move faster than spherical particles in mucosal environments.^13,41^ Yu *et al*. attributed this phenomenon to the rotational dynamics of nanoparticles facilitated by mucin fibers and shear flow within mucus.^41^ By contrast, Bao *et al*. demonstrated that rods of the same length, but differing in flexibility, can exhibit significantly different diffusion coefficients; flexible rods diffused faster than spheres, while rigid rods moved slower.^13^

### Diffusion dynamics of ligand-functionalized DNA origami

After characterizing the diffusion behaviour of naked DNA origami, we aimed at investigating the effect of ligand functionalization. We selected the anti-EGFR antibody as a functional cargo due to its relevance in colorectal cancer therapeutics, where its delivery across mucus is highly desirable.^28^ To this end, Anti-EGFR was conjugated to a ssDNA oligonucleotide using a click-chemistry approach. First, a dibenzocyclooctyne (DBCO) group was coupled to the antibody via an NHS-mediated coupling reaction, which was subsequently conjugated to a N_3_-modified ssDNA oligonucleotide. The resulting antibody-DNA conjugates were then incubated with the DNA origami nanostructures displaying complementary ssDNA sequences on their surface (**Figure 3A**). Gel electrophoresis and AFM analysis confirmed the successful conjugation of antibodies onto the DNA origami surface (**Figure SI 2**). Once assembled, DNA origami-antibody conjugates were tracked in mucus and analyzed as previously described. Surprisingly, we observed that the rods conjugated with the antibody exhibited a significantly higher MSD slope (**Figure 3B**), compared to the unconjugated nanorods, indicating an enhanced diffusivity, which can be observed in **Figure 3C**, showing the diffusion coefficient values for the different samples. For DNA origami rods, the enhancement in diffusion was accompanied by a significant increase in the α value, suggesting a shift from subdiffusive behavior to Brownian diffusion. In contrast, for the rectangles and icosahedra, the opposite behavior was observed. These shapes displayed a decrease in their diffusivity, as indicated by a lower MSD and diffusion coefficient compared to the naked DNA origami structures, and the decrease in their α values suggested a shift from diffusive to subdiffusive behavior. This effect could be due to the geometric differences in the structures, leading to different degrees of anisotropy. The antibody used in our study is a positively charged protein, and previous research has shown that the asymmetric charge distribution and spatial configuration of charges on the surface of the carrier can significantly impact the rate of transport, enhancing their diffusivity in mucus^42,43^ This suggests that both the antibody charge and its spatial arrangement on the DNA origami rods may play a crucial role in enhancing the diffusivity of the nanostructures, which is not observed for the other shapes. To confirm this hypothesis, we performed an additional experiment where we substituted the antibodies by Bovine Serum Albumin (negatively charged). Here, we observed that BSA conjugation also induced an increased diffusivity, though it was significantly less pronounced than with Anti-EGFR. This suggests that enhanced diffusivity in mucus requires an assymetric charge distribution (**Figure SI 3**). We conducted an additional control experiment where we substituted mucus by glycerol. Interestingly, we observed that the diffusivity of antibody-conjugated nanorods was not significantly enhanced compared to naked nanorods (**Figure SI 4B)** However, this behavior differs in mucus, where factors such as shape and antibody distribution appear to play a more significant role in motility. This suggests that while size influences diffusion in simpler media, the complexity of mucus requires consideration of additional parameters, such as surface properties and the spatial distribution of functional groups, to fully understand the particles’ movement. Therefore, we hypothesize that introducing asymmetric charge surfaces and carefully arranging these asymmetric charges can enhance particle mobility in mucus. This is further confirmed by the notion that viruses, which present both positive and negative charges on their surfaces, may owe their rapid movement through mucus to the specific spatial arrangement of charges.^42,44^

**Figure 3.**
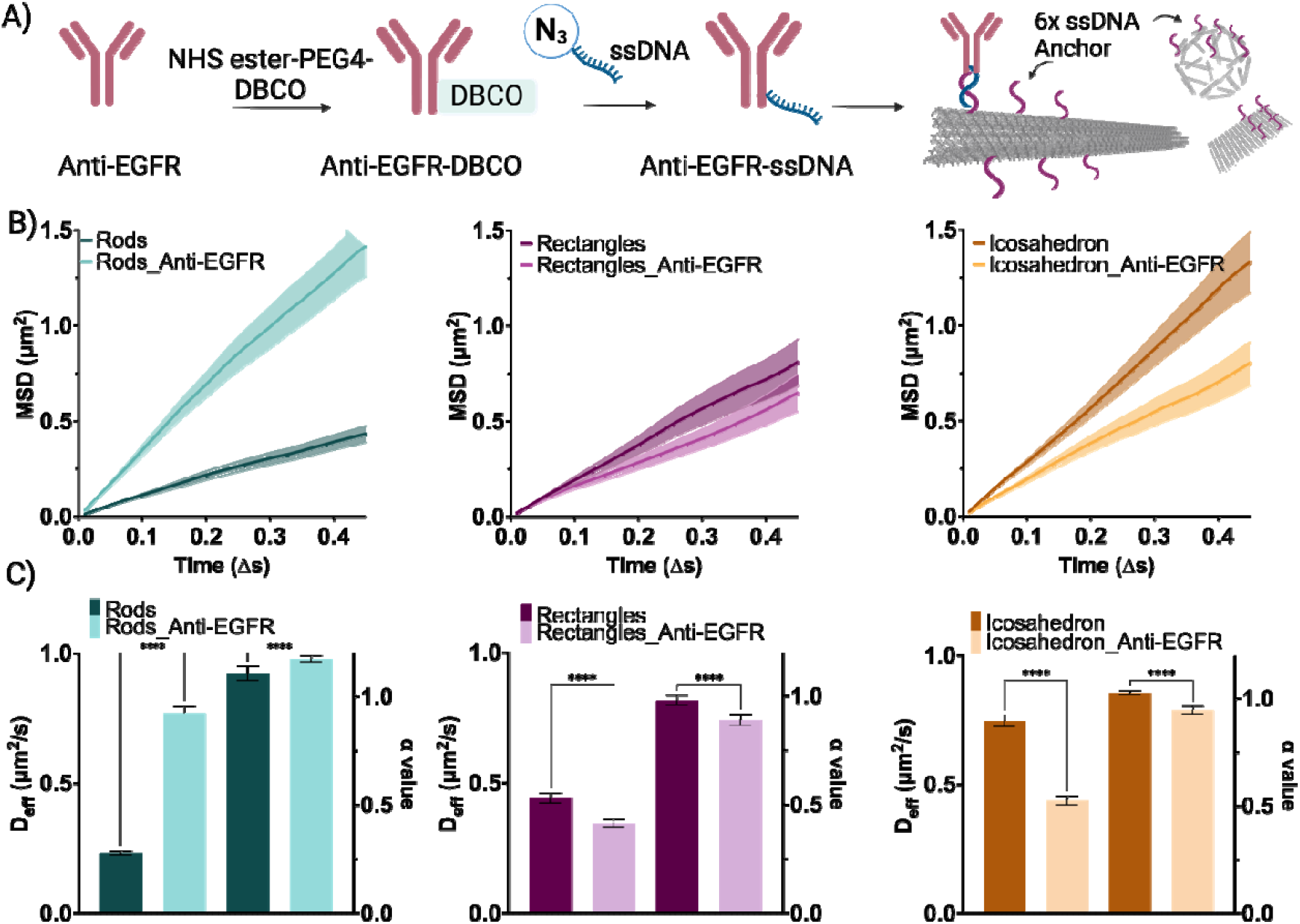
DNA origami conjugation. **A)** Illustrative diagrams depicting the functionalization of proteins via an NHS-mediated coupling reaction, followed by click chemistry with the azide-modified complementary handle on the DNA origami, and finally incubation with the DNA origami using a modified extended staple. **B)** Mean square displacement (MSD) analysis for different naked shapes and anti-EGFR conjugated DNA origami. **C)** Diffusion coefficients and alpha values extracted from the MSD analysis for the various shapes conjugated with antibody. Results are shown as the mean ± standard error of the mean (N=50). Asterisks ‘****’ denote statistically significant differences between groups, with p ≤ 0.0001.

### Ligand display effects on DNA origami diffusion across mucus

To further investigate the effect of antibody functionalization on diffusion, we designed DNA origami shapes with 0, 3, 6 and 9 antibodies (**Figure 4A**) and we monitored their diffusion acros mucus and glycerol. AFM and gel electrophoresis confirmed the successful conjugation of varying densities of anti-EGFR to the DNA origami (**Figure 4B**). In glycerol, as expected, we observed an inversely proportional effect, where the diffusion coefficient decreased at increasing antibody numbers functionalized on the DNA origami nanostructures. We attribute this to the increase in hydrodynamic size, which results in lower diffusivity (**Figure SI 4C**). By contrast, in mucus, we did not observe a consistent trend among shapes (**Figure 4C**). Our observations revealed that for modified rods, the highest diffusion coefficient was achieved with six antibodies conjugated to the surface. These results support our hypothesis that not only the charge but their distribution on the nanoparticle surface is a key determinant for their diffusivity across mucus, and that this is dependent on the nanoparticle shape. In the case of 6x functionalized rods, we assume that this is the most anisotropic charge distribution, and therefore the highest diffusivity is observed. By contrast, for rectangles, 3x antibody functionalization slightly increased their diffusivity, but at increasing antibody number, diffusivity decreased, reaching values even lower than the control, non-functionalized rectangles. In the case of icosahedrons, antibody functionalization did not only improve diffusivity but resulted in a significant reduction in all cases. In previous reports, transient, low-affinity interactions that allow particles to bind and release from mucin dynamically have previously been shown to increase the diffusivity of nanoparticles in mucus.^45,46^ Our results align with these observations, where functionalization with low or medium density of positively charged antibodies may lead to dynamic, low affinity interactions with negatively charged mucin, whereas the functionalization with higher antibody density results in strong electrostatic interactions with negatively charged mucin.

**Figure 4.**
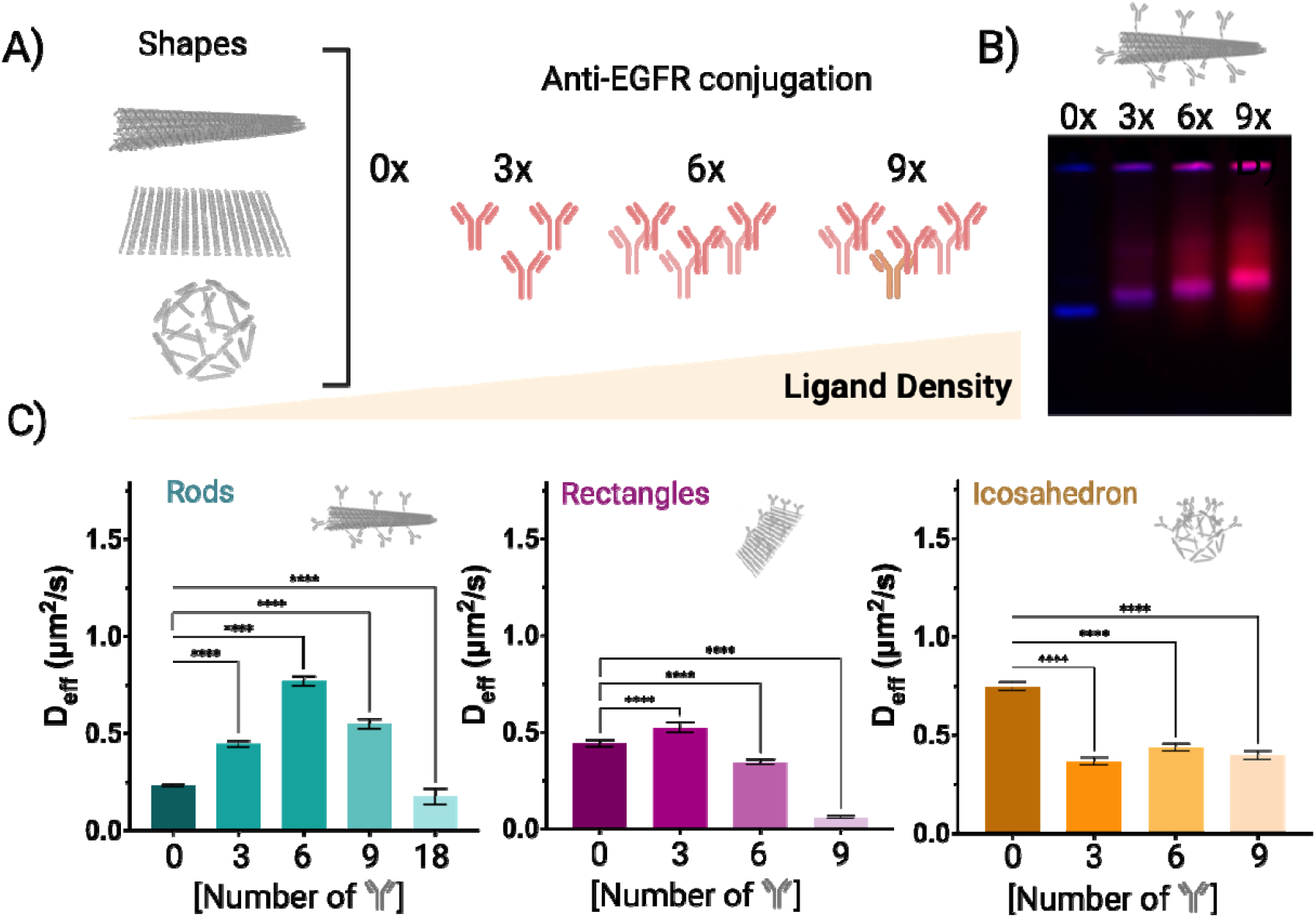
DNA origami density effect. **A)** Illustrative diagrams depicting the functionalization of at different densities on the DNA origami. **B)** Gel electrophoresis results of I) DNA origami structure; II) DNA origami structure-3x aEGFR; III) DNA origami structure-6x aEGFR; IV) DNA origami structure-9x aEGFR. **C)** Diffusion coefficients extracted from the MSD analysi for the different densities of antibody conjugated to various shapes. Results are shown as the mean ± standard error of the mean (N=50). Asterisks ‘****’ denote statistically significant differences between groups, with p ≤ 0.0001.

Overall, our results indicate that there is a threshold where DNA origami functionalization with positively charged antibodies does not increase diffusivity. These findings underscore the critical role of nanoparticle topological characteristics, including ligand charge, density, and particle shape, in governing mucus transport. A comprehensive understanding and precise control of these parameters will be essential for the rational design and optimization of nanoparticle-based drug delivery systems aimed at traversing mucosal barriers effectively.

## Conclusions

This work establishes DNA origami as a versatile and precisely tunable platform to decouple and systematically investigate the effects of nanoparticle topology (specifically shape, ligand functionalization and ligand density) on mucus transport, while keeping material composition constant. Through high-resolution single-particle tracking, we captured the spatiotemporal dynamics of particle motion, yielding mechanistic insights into how anisotropy and surface presentation modulate nanoparticle mobility through complex mucosal environments. Among the geometries studied, unmodified rod-shaped DNA origami exhibited the lowest diffusion coefficients; however, upon surface functionalization with a therapeutic antibody, their diffusivity increased markedly, surpassing that of both rectangle-shaped and icosahedral counterparts. This enhancement was maximized at an intermediate ligand density (six antibodies per particle), suggesting a geometry-specific optimum in surface presentation. In contrast, rectangles and icosahedra showed only modest increases in diffusivity at low ligand densities (three antibodies per particle), with diminished transport at higher functionalization levels. These results highlight the nuanced, non-monotonic relationship between ligand density and particle mobility and underscore the importance of shape-dependent optimization in the design of mucus-penetrating delivery systems. Collectively, our findings establish DNA origami as a powerful tool for elucidating transport mechanisms and guiding the rational design of next-generation nanocarriers for mucosal drug delivery.

## CRediT authorship contribution statement

**Matteo Tollemeto** (Conceptualization; Data curation; Formal analysis; Investigation; Methodology; Validation; Visualization; Writing – original draft; Writing – review & editing) **Lars J.M.M. Paffen** (Conceptualization; Formal analysis; Investigation; Methodology; Validation; Writing – review & editing) **Lasse Højlund Eklund Thamdrup** (Conceptualization; Supervision; Writing – review & editing) **Anja Boisen** (Conceptualization; Supervision; Visualization; Writing – review & editing; Project administration; Funding acquisition) **Jan van Hest** (Conceptualization; Supervision; Visualization; Writing – review & editing; Project administration; Funding acquisition) **Tania Patiño Padial** (Conceptualization; Supervision; Visualization; Writing – original draft; Writing – review & editing; Funding acquisition)

## Acknowledgements

The authors would like to acknowledge the Danish National Research Foundation (DNRF122) and Villum Fonden (Grant No. 9301) for intelligent drug delivery and sensing using microcontainers and nanomechanics (IDUN) and the Novo Nordisk Foundation (NNF17OC0026910). T.P. thanks the Irene Curie fellowship and the ICMS.

## Conflict of Interest

The authors declare no conflict of interest.

## Notes

### Competing Interest Statement

The authors have declared no competing interest.

