## Supporting Information for "Topology Determines DNA Origami Diffusion in Intestinal Mucus"

**Corresponding authors:**

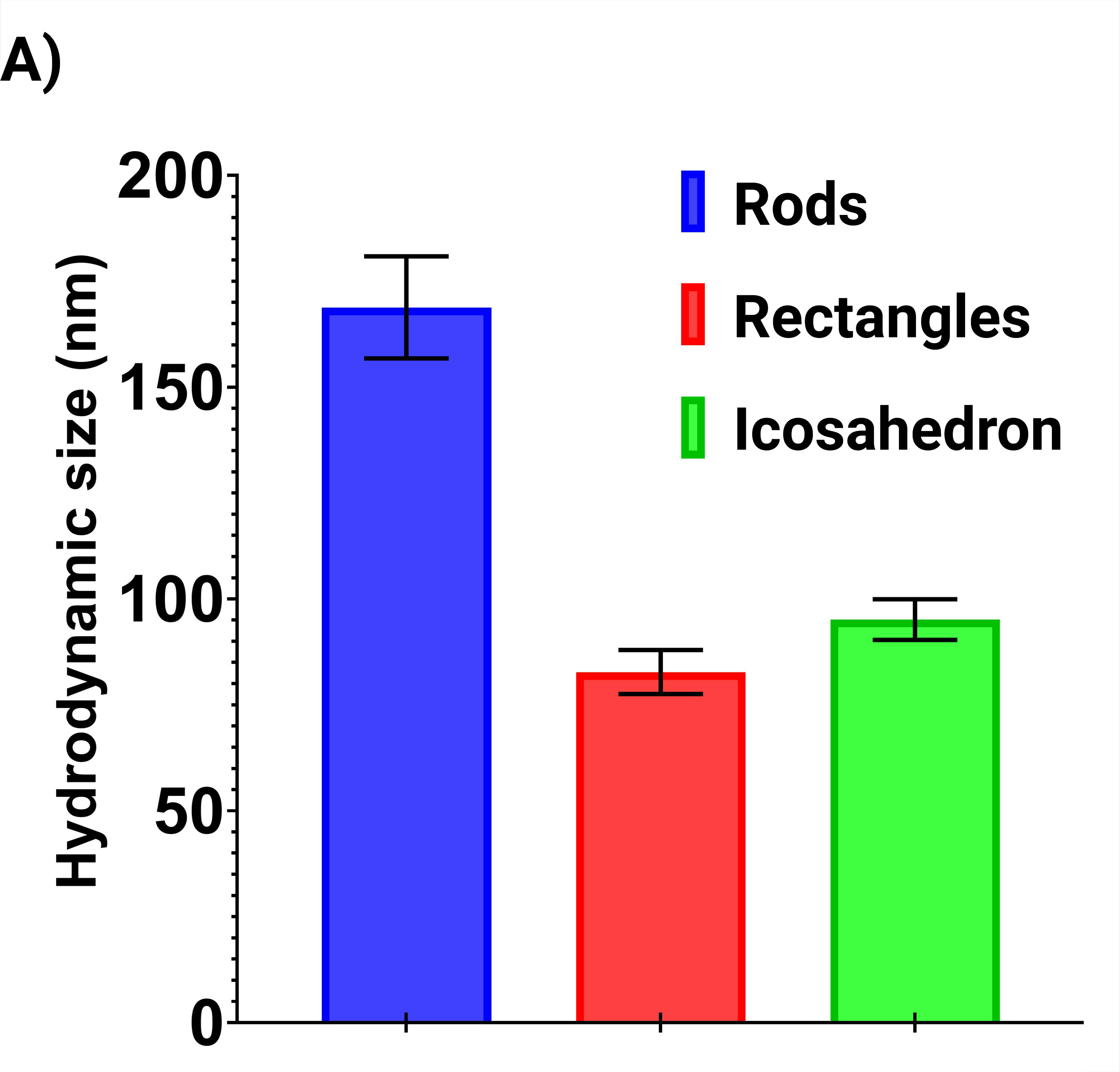

**Figure SI 1|** A) DLS analysis shows the varying hydrodynamic radii of the different DNA origami structures.

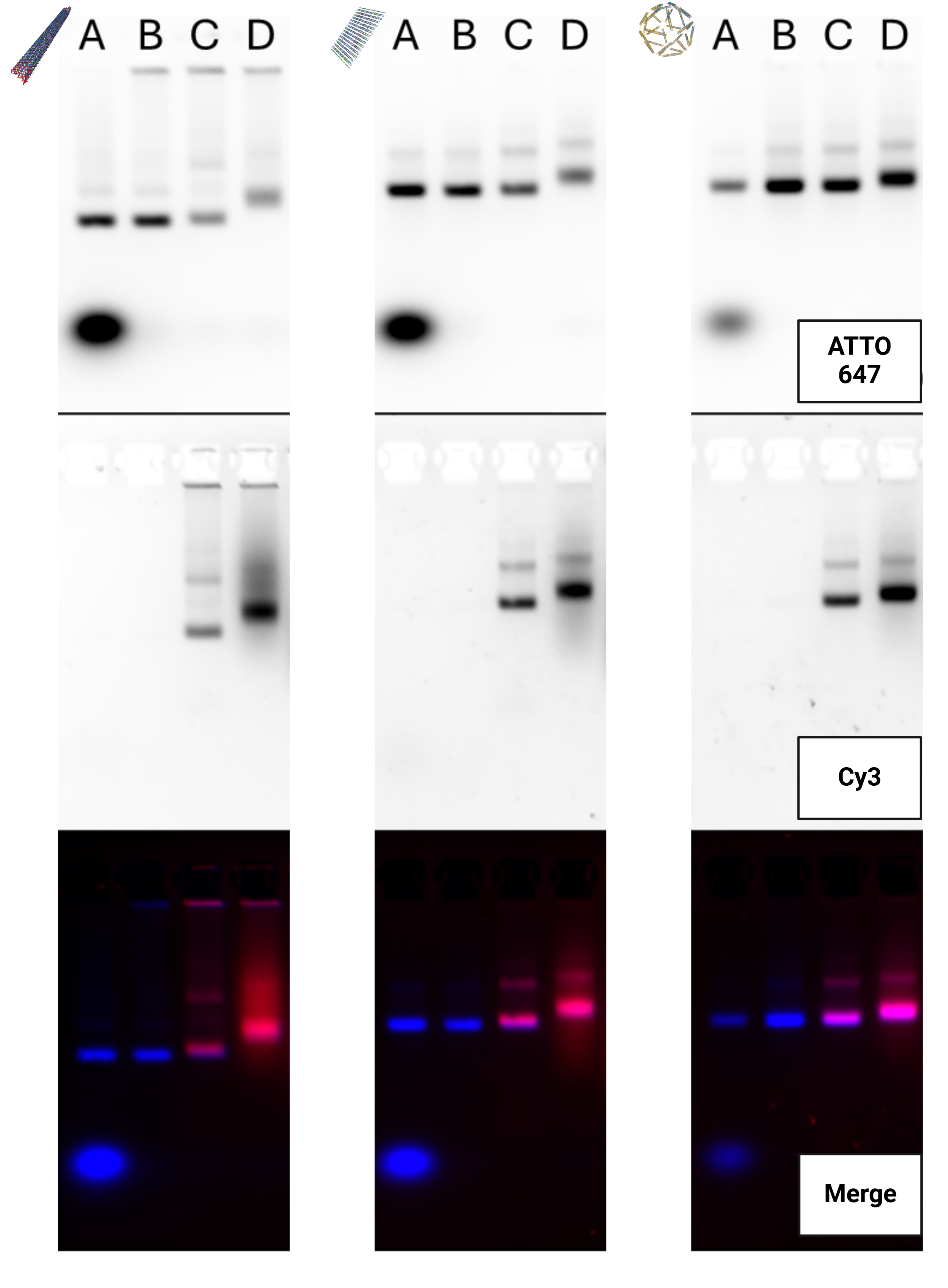

**Figure SI 2|** Gel electrophoresis results of A) DNA origami structure (non-purified); B) DNA origami structure (purified); C) DNA origami structure-BSA; D) DNA origami structure-aEGFR;

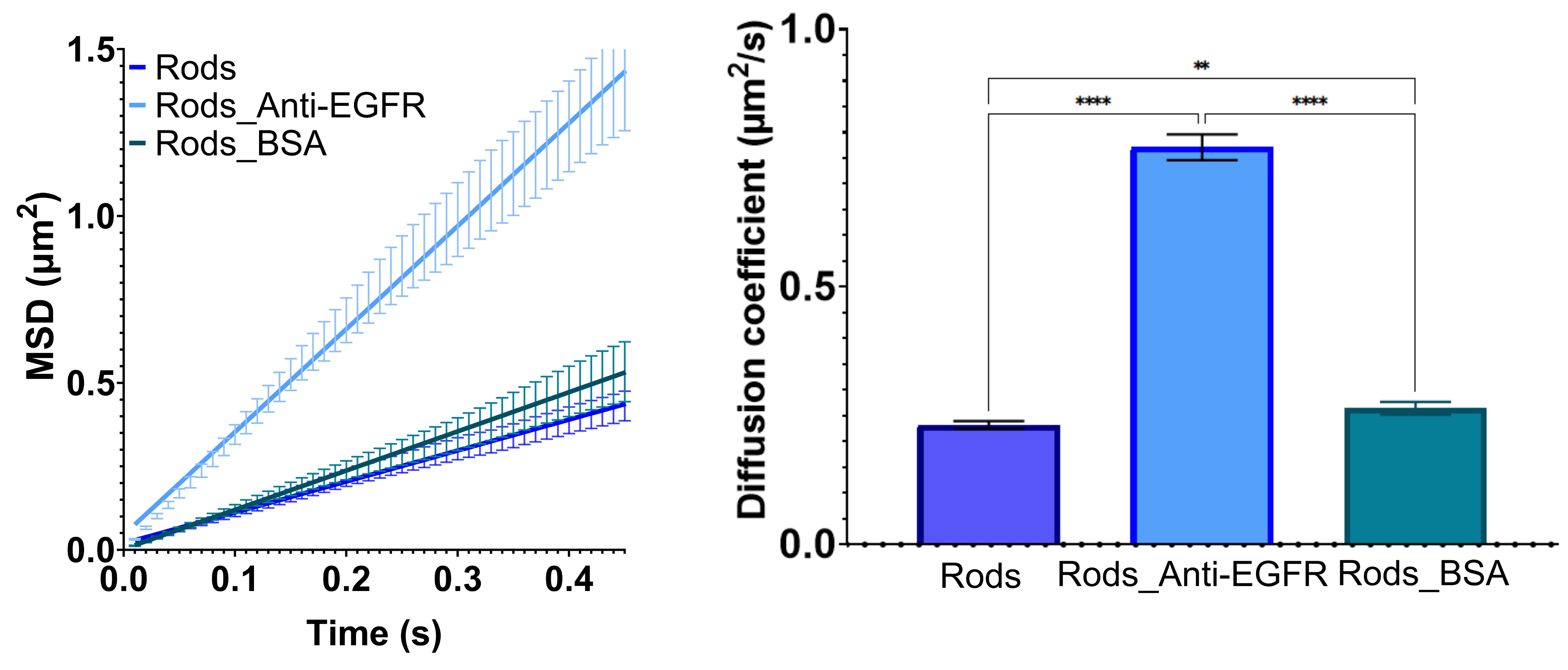

**Figure SI 3|** MSD and diffusion coefficients for DNA origami structure; I) DNA origami structure; II) DNA origami structure-aEGFR; III) DNA origami structure-BSA.

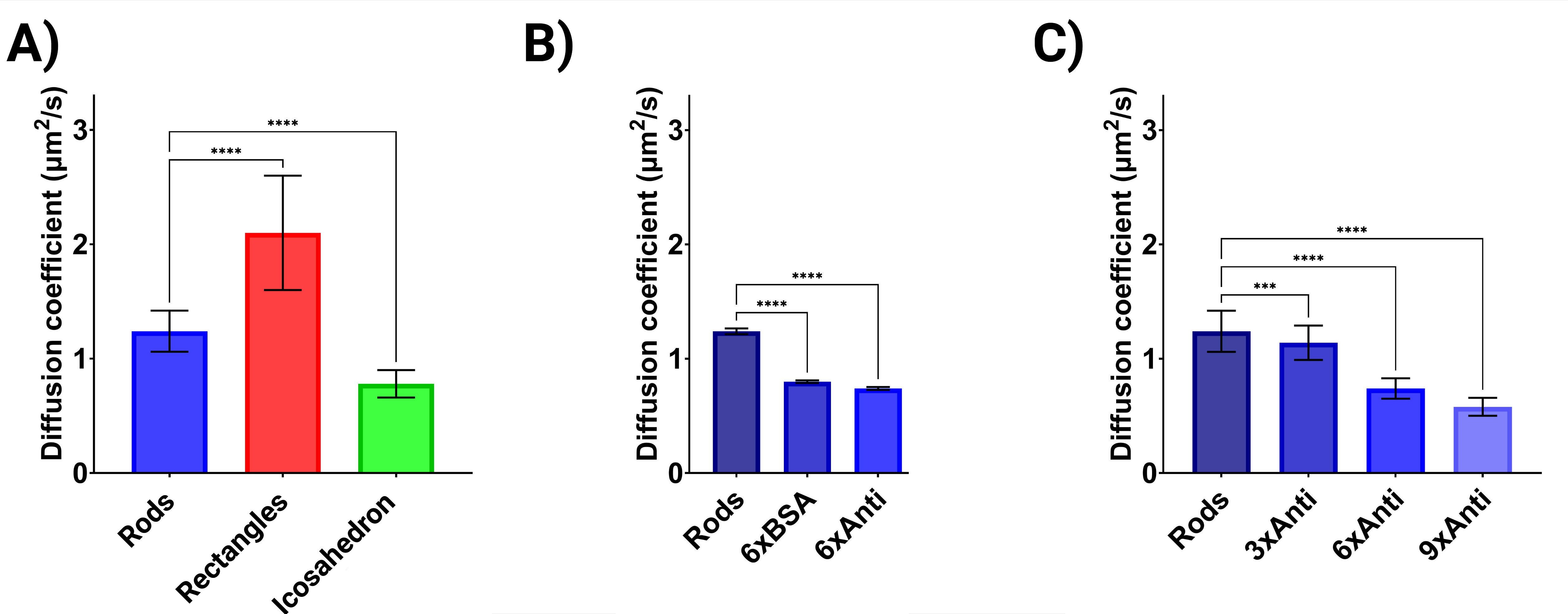

**Figure SI 4|** Diffusion coefficients derived from MSD analysis in glycerol 50% for: A) Different DNA origami shapes; B) Various protein conjugations; C) Varying densities of antibody conjugation to the rods.

**Table SI 1|** Theoretical Calculations of Surface Area and Volume values.

|  | **Rods** | **Rectangles** | **Icosahedron** |
| --- | --- | --- | --- |
| **Surface Area (nm^2^)** | **10436** | **12984** | **7853** |
| **Volume (nm^3^)** | **53218** | **11718** | **65449** |
| **Surface Area/Volume** | **0.19** | **1.10** | **0.12** |

**Materials and Methods**

**Materials**
All chemicals were used as received unless otherwise stated. Staple strands, handle-extended staple strands, modified DNA oligonucleotides, and fluorescently modified DNA oligonucleotides, the p7560 scaffold for the rods and M13mp18 scaffold for the rectangles and icosahedron were obtained from Integrated DNA Technologies. InVivoMAb anti-human EGFR was purchased from Bio X Cell. All other chemicals and reagents, including: BSA, catalase and urease polyethylene glycol, phosphate-buffered saline (PBS), dimethyl sulfoxide (DMSO), glycerol, toluene, NaCl, MgCl_2_, EDTA, RNase-free MQ were purchased from Sigma-Aldrich (St. Louis, MO). Solvents were HPLC grade and used as received. Ultrapure water was used throughout the experiments and obtained from Merck Millipore Q-Pod system (Merck Group, Burlington (MA), U.S.A) with a 0.22 μm Millipore Express 40 filter (18.2 MΩ).

**Data and statistical analysis**

Data analysis was performed with GraphPad Prism (Version 9.4.1 (681) Insight Partners, Graphpad Holdings, LLC, New York City (NY), U.S.A.). Data are presented as average and standard derivation (SD) where n represents the number of repetitions within each sample and N represents the number of samples. Statistical analysis was performed for statistically significant differences (*p < 0.05) with GraphPad Prism using a t-test (for two independent populations) or one-way analysis of variance (ANOVA) (three or more independent populations). The figures were created with Biorender.com. Image analysis and processing was done using the software ImageJ (version 1.53t, National Institutes of Health, USA).

**DNA origami assembly and purification.**
The folding reaction was carried out by first thawing all components, on ice for 20 minutes. To prepare the various buffers used in the experiment, several stock solutions and mixtures were created. For the 10X Pre-Fold/Precipitation buffer, 500 µL of 1 M Tris and 200 µL of 0.5 M EDTA were combined with 9.3 mL of RNase-free MQ, achieving a final buffer composition of 50 mM Tris and 10 mM EDTA at pH 8.5. The folding buffer was prepared by mixing 1200 µL of 100 mM MgCl_2_, 500 µL of 500 mM NaCl, 1000 µL of the 10X Pre-Fold/Precipitation buffer, and 7.3 mL of RNase-free MQ, resulting in a final composition of 5 mM Tris, 25 mM NaCl, 12 mM MgCl_2_, and 1 mM EDTA at pH 8.5. Finally, the storage buffer was made by dissolving 2 PBS tablets in 200 mL of MQ to create a 2X PBS stock, which was filter sterilized. A 1 M MgCl_2_ solution was prepared by dissolving 50.8 g of MgCl_2_ in 250 mL of MQ. The storage buffer was then prepared by mixing 100 µL of 1 M MgCl_2_, 5 mL of 2X PBS stock, and 4.9 mL of RNase-free MQ, resulting in a final composition of 1X PBS and 10 mM MgCl_2_ at pH 7.4. The tubes were then centrifuged at 500×g for 1 minute.

A typical workflow (**Table SI 3**) for the scaffold (P7560 for the rods and M13mp18 for the rectangle and icosahedron), staple strands and buffers is shown below. The mixture was then thoroughly mixed by pipetting and gently inverting the tube. Finally, the total reaction volume was aliquoted into 200 μl PCR tubes, with each tube containing 50 μl of the mixture. All steps were performed using RNase-free equipment to maintain sample integrity.

To conduct the PCR reaction, the thermocycler was programmed with the following parameters for the rods: an initial denaturation step at 80°C for 15 minutes, followed by a gradient annealing phase starting at 79°C, held for 1 minute, and repeated 19 times. This was succeeded by a gradient extension phase beginning at 59°C, held for 23 minutes and 20 seconds, and repeated 35 times. The final step involved holding the samples at 4°C indefinitely. The PCR reaction for the rectangles and icosahedron was as followed: mixture heated to 95 °C for 15 min and then slowly cooled to 20 °C at a rate of 1 °C/min.

To purify the origami product using PEG precipitation, a PEG buffer was first prepared by combining 500 µL of 2X precipitation buffer, 300 µL of PEG 8000 (50% w/v), and 200 µL of RNase-free molecular-grade water, resulting in a final concentration of 15% (w/v) PEG. The PEG buffer was added to the reaction mixture in a 1:1 (v/v) ratio. The mixtures were then gently mixed by pipetting and tube inversion, followed by incubation on ice for 10 minutes. After incubation, the tubes were centrifuged at 21,000×g and 16°C for 25–45 minutes. The supernatant was carefully removed using a pipette, and the resulting pellet was redissolved in 25 µL of storage buffer. The samples were incubated at 30°C for 30 minutes to dissolve the pellet, followed by gentle pipetting and an additional 10 minutes at 30°C.

DNA origami concentration was measured using a Nanodrop ND-1000 UV-Vis Spectrophotometer (NanoDrop Technologies, Inc., Wilmington, DE, USA). 1 µL of the sample mixture was added and absorption at 260 nm was measured, using an extinction coefficient (ε) of 9.45 × 10^7^ M^-1^*cm^-1^ (rods), 1.24 x 10^8^ M^-1^*cm^-1^ (rectangles), 9.45 x 10^7^ M^-1^*cm^-1^ (icosahedron). The path length (l) of the measurement was set to 10 mm. The concentration of the DNA in the sample was calculated using the Lambert-Beer law, equation (1) as follows:

$$\boldsymbol{c:}\frac{\boldsymbol{A}_{\boldsymbol{260}}}{\boldsymbol{\varepsilon x l}} \left[ \boldsymbol{M} \right]$$

To confirm DNA origami self-assembly, agarose gel electrophoresis was conducted. A 1 L agarose gel running buffer was prepared using 0.5x TBE and 10 mM MgCl2, adjusted to pH 8.0. The gel tray was prepared by sealing the open ends with autoclave tape to prevent leaks. A 0.6% agarose gel was made by weighing 600 mg of agarose into a 100 mL Erlenmeyer flask, adding 40 mL of the prepared running buffer, and heating the mixture in a microwave until the agarose fully dissolved. Once dissolved, 4 μL of 10,000X SYBR SAFE DNA staining dye was added to the agarose solution, which was mixed well by swirling. The mixture was poured into the tray, allowed to solidify, and a comb with small teeth was placed at one end to create wells. Samples for electrophoresis were prepared by mixing the origami with a loading dye made from 7.5% (w/v) Ficoll-400 dissolved in MQ. After the gel solidified, the tape was removed, and the gel was placed in the running chamber with the well side facing the black electrode. The chamber was filled with running buffer until the gel was submerged, and the comb was carefully removed. Samples (10 μL each) were loaded into the wells, and the gel was run at 65 V for 90 minutes, with ice placed in the bucket containing the running chamber to maintain a low temperature. After electrophoresis, the gel was imaged to analyze the results.

**Protein conjugation to an ssDNA handle.**
Proteins were first conjugated to DBCO-PEG4-NHS (8 molar equivalent to the protein) and NHS-Cy3 (8 molar equivalent to the protein). Each protein was initially dissolved in PBS, while DBCO-PEG4-NHS and NHS-Cy3 were dissolved in DMSO. The three reagents were mixed in a 1:10 (v:v, DMSO:PBS) solution and incubated at 300 rpm for 6h at RT. Following incubation, the conjugated protein was purified through overnight dialysis against PBS using Spectra/Por dialysis membrane tubing with a 12-14 kDa molecular weight cutoff. The dialyzed mixture was then concentrated using a 100 kDa MWCO Amicon spin filter, involving multiple wash steps with PBS to remove unbound components. The protein concentration and labeling efficiency were determined by measuring the absorbance at 280 nm for protein content and at 309 nm and 535 nm for DBCO and Cy3, respectively.
To couple the protein-DBCO-Cy3 protein conjugate to ssDNA, it was mixed with Seq1-N_3_ (20 molar equivalent to the protein) and the mixture was incubated overnight at 4°C. Post-incubation, excess unbound oligonucleotides was removed using spin filtration with an Amicon 100 kDa MWCO spin filter 4 times. Finally, the labeled protein-oligonucleotide conjugate was recovered by performing a reverse spin at 1,000 × g for 1 minute and then diluted back to the original volume of 100 μL using PBS. The successful labeling of the oligonucleotide was confirmed by SDS-PAGE analysis.

**Antibody complexation to DNA origami.**
The assembly process between the nanorods and ligands was carried out over a 3-hour period, starting with 1 hour at 37°C followed by 2 hours at 22°C. During this reaction, a threefold molar excess of protein was used for each nanorod handle (3, 6 and 9), a typical workflow is shown below (**Table SI 4**). This allowed the handles on the origami structures to interact with complementary sequences on the protein-ligands, forming double-helix DNA structures.

**DNA origami characterization – Atomic Force Microscopy**

Topographic images were captured in tapping mode under liquid conditions using a Cypher ES Environmental atomic force microscope (Oxford Instruments). Cantilevers with sharpened silicon tetrahedral tips, a nominal spring constant of 0.09 N/m, and a peak resonance frequency in water around 25 kHz (BL-AC40TS, Olympus) were utilized. Substrates for sample application were prepared by attaching round laser-cut mica discs (ca. 1 cm², Ted Pella) to metal sample discs using double-sided adhesive tabs. DNA origami solutions were diluted to 2 nM in Imaging buffer (1x TAE, 1 mM EDTA, 10 mM MgCl2, pH 8.0), and 5 µL of this solution was deposited onto the freshly cleaved mica disc. After a 30-second incubation, 50 µL of Imaging buffer was added on top. Before measuring, 80 μl of Imaging buffer water was applied to the cantilever and the cantilever was manually lowered until the droplets on both the cantilever and the mica fused. The correction collar on the objective was set to 2.0. Images (512×512 px) were acquired over areas varying from 5.0 µm² to 0.5 µm², with scanning and feedback parameters optimized for each image. All images were analyzed using Gwyddion v2.64.

**Size and Surface Charge**

The hydrodynamic size, polydispersity index, and ζ-potentials of the DNA origami were measured at 37°C using a Malvern Zetasizer Nano ZSP (Malvern Instruments, Worcestershire, U.K.) in single-use disposable microcuvettes. Measurements were conducted at a DNA origami concentration of 10 nM. Dynamic light scattering was performed on 100 μL samples at a detection angle of 173°, with 13 runs of 10 seconds each, across three measurements. The ζ-potential was assessed using laser Doppler electrophoresis with the Zetasizer Nano ZS (Malvern Panalytical). For this, 700 μL of the sample was measured in folded capillary cells, with 10 runs per measurement, repeated three times. (N=3, n=3).

**Mucus isolation** 
Intestines from 3 healthy fasted (18–24 h) piglets were obtained from LifeTec Group. Immediately after euthanization, up to 5 m jejunum was isolated distal to the ligament of Treitz. Sections were opened by a latitude cut and porcine intestinal mucus was isolated by gently scraping the mucosal surface. Mucus was kept on ice at all times and stored at −20 °C until use.

**DNA origami tracking experiment in mucus**

The DNA origami was mixed with mucus at a 10:1 ratio (v/v, mucus:DNA origami), resulting in a final DNA origami concentration of 1 nM. The mixture was then incubated at 37°C. Similarly, this was repeated for the measurements in glycerol 50%.

**DNA origami tracking acquisition and analysis**

Images were acquired in a Nanoimager® (ONI, Oxford) using the NimOS software. ATTO-647 labelled origamis were imaged with a 640 nm laser (190 mW). The sample was illuminated using a highly inclined and laminated optical sheet (HiLo) at an angle of 47°, at 10% laser power. Fluorescence was recorded using an ONI 100×, 1.49 NA oil immersion objective and passed through a quad‐band pass dichroic filter. Images were acquired onto a 425×518 pixel region (pixel size 0.117 μm) for 10 s at 100 fps. For each experimental condition, a total of at least 3 different biological samples were analyzed.
The results were filtered using the NimOS software with the following settings: maximum frame gap = 5, minimum distance between frames = 0.800 µm, exclusion radius = 1.200 µm, a minimum number of steps = 50, a minimum diffusion coefficient of 0.01 μm²s⁻¹. With these parameters, track steps were extracted, and analyzed using a Python-based code^1^ to obtain the trajectories of the DNA origami (n = 50) and calculate the mean-squared displacement (MSD) with the following equation (2):

$$\boldsymbol{MSD=<}\boldsymbol{r}^{\boldsymbol{2}}\left( \boldsymbol{t} \right)\boldsymbol{> =<(}\frac{\boldsymbol{1}}{\boldsymbol{N}}\sum_{\boldsymbol{i=0}}^{\boldsymbol{N}} \left( \boldsymbol{r}_{\boldsymbol{i}}\left( \boldsymbol{t} \right)\boldsymbol{-}\boldsymbol{r}_{\boldsymbol{i}}\left( \boldsymbol{0} \right) \right)^{\boldsymbol{2}}\boldsymbol{>}$$

Then, the diffusion coefficient was obtained by fitting the MSD data to equation (3), where r = radius and t = sampling time and MSD(t) = 2dD, where D = diffusion coefficient and d = dimensionality (ONI measurements have dimension d = 2):

$$\boldsymbol{MSD=}\left( \boldsymbol{4}\boldsymbol{D} \right)\boldsymbol{\Delta t}$$

was used to fit the MSD curves.

$$D=\frac{k_{B}T}{6\pi\eta R_{h}}$$

where *D* = diffusion coefficient, k_B_ = Boltzmann constant, *T* = temperature, *η* = mucus viscosity

and Rh = hydrodynamic radius.
